# Cognition and behaviour in learning difficulties and ADHD: A dimensional approach

**DOI:** 10.1101/260265

**Authors:** Susan E. Gathercole, Duncan A. Astle, Tom Manly, the CALM Team, Joni Holmes

## Abstract

**Background:** Academic underachievement often accompanies the symptoms of inattention and hyperactivity/ impulsivity associated with ADHD. The aim of the present study is to establish whether learning difficulties have the same cognitive origins in this comorbid condition as in children who do not have ADHD.

**Methods:** Participants were 163 school-aged children with learning difficulties. Over a third also had a diagnosis of ADHD. Cognition, behaviour and learning attainments were assessed.

**Results:** The sample was distinguished by three cognitive and three behavioural dimensions. Learning was equivalently related to cognitive dimensions for children with and without ADHD. A diagnosis of ADHD was associated only with elevated levels of ADHD symptoms and problems with emotional control.

**Conclusions:** Distinct dimensions underpin academic learning and the control of impulsive and emotional behaviour impaired in ADHD. Phonological deficits are associated with learning problems in literacy and maths, and impairments in nonverbal and executive abilities with mathematical learning difficulties. The comorbid condition of ADHD combined with learning difficulties reflects independent deficits in the cognitive dimensions critical for learning and in the control of impulsive and emotional behaviour.

---

Learning difficulties take many different forms. Children can fail to learn to read and write at the expected rate, to understand written language, or to grasp numerical concepts and develop reasonable competence in arithmetic and more complex areas of mathematics. Many children with these learning difficulties also meet diagnostic criteria for ADHD, a psychiatric diagnosis based on observation of behaviours such as apparent inattention, difficulties waiting one’ s turn, poor organisation and excessive restlessness that are considered anomalous and occur in at least two settings (APA, 2013). For example, in a US population study, 44% of children with ADHD also had learning difficulties and 42% of those with a specific learning disability also met criteria for ADHD (Pastor & Reuben, 2008), an association well above chance levels.

The present study aimed to identify whether cognitive and behavioural dimensions could distinguish children who have both learning difficulties and ADHD from those that have learning problems alone. Four broad types of model could apply. The first is that ADHD carries with it a vulnerability to cognitive impairments that, as with the non-ADHD population, are risk factors for particular learning problems. Reading problems have been linked with slow processing speed (Willcutt et al., 2010), poor phonological skills (Melby-Lervag, Lyster, & Hulme, 2012), and weak short-term memory (STM) and WM for verbal material (Ramus, Marshall, Rosen, & van der Lely, 2013; Swanson & Sachse-Lee, 2001). Maths difficulties, in contrast, are more closely linked with impairments in visuo-spatial working memory (Szucs, Devine, Soltesz, Nobes, & Gabriel, 2013) as well as executive functions (EFs) including poor inhibitory control and planning (Bull & Scerif, 2001). If this vulnerability is independent of the level of ADHD symptoms, we would expect comorbid children to resemble non-ADHD children with learning difficulties in terms of their cognitive profiles and to differ only with respect to the higher frequency of ADHD behaviours in the former group. The frequency of ADHD behaviours would be independent of learning outcome.

The second broad possibility is that the features that give rise to the ADHD diagnosis and that distinguish these children from others with learning difficulties are linked with particular learning outcomes in a manner that is relatively independent of cognitive capacity. It could be, for, example, that some areas of learning (e.g. maths) are particularly sensitive to fluctuating attention or an impulsive urge to complete and move on. In this case we might expect children with and without ADHD, but with the same level of attainment, to have rather different cognitive profiles. Here the frequency/severity of ADHD behaviours would be related to learning outcomes only in the comorbid group.

A third possibility is that children with ADHD have a combined risk and that both specific cognitive impairments and the disruptive influence of the ADHD symptoms will affect learning outcomes. Compared to children with learning difficulties but no ADHD diagnosis, these children would therefore be expected to have similar levels of cognitive impairment but disproportionately poor learning outcomes. ADHD behaviour ratings would be related to attainment scores but independently of cognitive skills.

Finally, ADHD may be linked with cognitive risk factors for poor learning that are, in turn, related to the severity of ADHD symptoms. We would therefore expect common cognitive profiles for both the children comorbid and specific learning difficulty children, and ADHD behaviour ratings that correlate with cognitive skills and attainment.

There are grounds to favour the first and third possibilities, both of which hold that cognitive impairment is a risk in ADHD which is relatively independent of the severity of the behaviours giving rise to the diagnosis (whether or not these behaviours pose an additional risk to school attainment). Holmes et al (2014) found that the cognitive profiles and academic achievements of children with low WM (and no ADHD) were largely indistinguishable from an ADHD sample, the groups differing only in impulsive and emotional behaviour. Furthermore, cognitive impairments in children with ADHD appear to be similar to those in children with reading difficulties (Cheung et al., 2012). However, other studies have reported direct links between behavioural symptoms specific to ADHD, cognition and reading abilities (Gremillion & Martel, 2012; McGrath et al., 2011).

Sonuga-Barke (2002; 2010) has conceptualised ADHD as a consequence of impairments in two separate neurobiological pathways. One serves executive functions such as working memory (WM), planning and switching. The other, less implicated in specific learning outcomes but contributing to hyperactivity and impulsivity, is involved in the regulation of motivation and more specifically, in processing the affective value of rewards. This distinction maps well onto a characterisation of cool executive functions (EFs) that control cognitive performance regardless of affective context, and hot EFs implicated in impulsivity and hyperactivity (Zelazo & Müller, 2002)). These two systems appear to be neurobiologically, functionally, and genetically distinct (Castellanos et al., 2005; Solanto, Arnsten, & Castellanos, 2001).

The approach we adopted to this issue in the present study was novel in two respects. Firstly, it used data-driven analyses to identify dimensions of cognition and behaviour rather than more conventional confirmatory model-testing methods. Studies of children with learning difficulties often base their interpretations of cognitive deficits on overly simplistic assumptions about individual tasks. For example, tasks such as nonword repetition, digit span and rhyme oddity detection have all been variously described as tests of phonological short-term memory, phonological sensitivity and the quality of phonological representations. But in reality, each of these tests is constrained by multiple skills, some shared and others that are task-specific (Gathercole, 2006). The present approach minimises such interpretational biases by identifying a small number of dimensions in a hypothesis-free manner.

A second novel feature of the study was its use of a large, heterogeneous sample of struggling learners recruited irrespective of diagnostic status or comorbidity. The children attended a research clinic for individuals with problems in attention, learning and memory, based on referrals by professionals working across a broad range of children’ s services in education and health. This sample is therefore highly representative of the population of struggling learners at large.

The approach used here echoes a broad shift in focus over the past decade away from diagnosis-specific deficits and symptoms towards dimensions of psychopathological disorders that cut across distinct disorders conventionally considered to be distinct. This has been reinforced by the priorities set for basic and translational research by the NIMH Research Domains Criteria project (Cuthbert & Insel, 2013). Although this approach has to date been most extensively applied to adult psychiatric conditions, there is now growing consensus that neurodevelopmental disorders too are better characterized by dimensions than diagnostic categories (Casey, Oliveri, & Insel, 2014; Sonuga-Barke & Coghill, 2014; Zhao & Castellanos, 2016). As yet, though, most studies still recruit single diagnostic groups with typical controls for comparison and employ the standard neuropsychological methods of hypothesis-driven analysis. In contrast the present study applies an exclusively data-driven approach in search of common and distinct dimensions of cognition and behaviour in a large mixed sample of children with learning difficulties and ADHD.

## Methods

### Study design

The Centre for Attention, Learning and Memory (CALM) is a research clinic established at the MRC Cognition and Brain Sciences Unit in Cambridge, UK. Referrals are sought from local specialist teachers, special educational needs coordinators, educational psychologists, clinical psychologists, child psychiatrists and ADHD nurse practitioners (through child and adolescent mental health services, CAMHS), and speech and language therapists. Our inclusion criteria were children aged 5-16 years in mainstream schooling considered to have problems in attention, learning, and/ or memory. Exclusion criteria were significant uncorrected problems of hearing or vision, a known genetic disorder, and not being a native English speaker. The route to referral was for the professional to give an information pack to the child’ s parents or carers who then may or may not have contacted CALM. Upon receipt of this expression of interest, a CALM team member contacted both referring agent and family member to check that the child met the study criteria. At this point, the referrer identified their primary reason for referral: problems in attention, literacy, maths, language or memory, or poor educational progress in general. If suitable the child and family were invited to attend an appointment in which the child completed a range of assessments and the parents/carers completed standardised as well as giving a family history. A report summarising the results of the measures was then returned to the referring professional.

### Ethical considerations

The study was approved by the local NHS Research Ethics Committee (13/EE/0157). Parents/ carers provided informed consent for participation in the study on behalf of the child.

### Participants

Recruitment to this study was based on the first 550 children attending the CALM clinic. Recruitment routes and selection criteria resulting in the sample of 163 children included in this study are shown in S1. Of the 217 children who remained in the sample after the exclusionary criteria above were applied, a further 54 children were excluded on the basis of academic achievement judged to be in the age-typical range, operationalized as a standard score greater than 85 in measures of reading, spelling, and maths. This cut-off score was selected in accordance with guidance for diagnosis of specific learning disorders (SLD) in the Diagnostic Statistical Manual of Mental Disorders 5 (APA, 2013), which recommends combining clinical judgment (satisfied by the specialist referrals) combined with scores below the 16% centile to indicate individuals who might have SLD. Children with intellectual disabilities (IQ scores 2 or more SDs on standardised assessment) are excluded from this diagnosis. In line with the inclusive dimensional approach adopted here, no exclusions were made on the basis of the IQ measure, matrix reasoning. Of the 163 children included in the present study, 15 (9.2%) had T-scores at or below 30, compared with 4.5% of children in the general population. Thus although the majority of the children are best characterised as having specific learning difficulties, the broader label of ‘learning difficulties’ is employed her to best characterise the sample.

The mean age of the sample was 10y 6m (range 8y 0m to 16y 1m) and there were 108 males (66%) and 55 females. Of these, 45 of the children (27%, 37m) had a diagnosis of ADHD. A further 17 (9%) children were under investigation for ADHD (13%, 14m). They were referred by a specialist ADHD nurse and were awaiting a full clinical assessment. At the time of reporting this process had not been completed. Twenty-six of 45 children diagnosed with ADHD had been prescribed psychostimulant medication. No information on medication status was recorded for the remaining 4 children with ADHD. Fourteen children were reported to have dyslexia, and 15 to have comorbid Autism Spectrum Disorder.

The primary reasons for referrals are shown in as a function of the service provider and ADHD group (ADHD, ADHD under investigation, or no ADHD) are shown in S3. The most common reasons for referral were problems in academic learning (35%) and in attention (37%). Attentional problems were identified as the primary reason for 80% of the children with ADHD, 82% of children under investigation for ADHD, and 10% of the children without ADHD. For the latter group, the most common primary reason was problems in academic learning (45%).

### Assessments

Each child completed assessments of cognition and learning administered by a team member in a quiet room in the CALM clinic. The carer completed a set of questionnaires relating to behaviour, communication, wellbeing and family history. Data are reported here for assessments of cognition and learning, and of behaviour relating to attention and EFs.

#### Phonological processing

The Alliteration and Rapid Automated Naming (RAN) subtests of the Phonological Abilities Test (Muter, Hulme, & Snowling, 1997) were administered. The RAN subtest required children to name aloud as quickly and accurately as possible object words. The Alliteration subtest assesses phonological awareness. Sets of three spoken words were read aloud and children were required to judge which two words started with the same sound. The number of correct responses formed raw scores, which were converted to standard scores (M = 100, SD = 15).

#### WM

Four subtests of the Automated Working Memory Assessment (Alloway, 2007) were administered. Standard scores (M= 100, SD=15) were calculated for each subtest. Digit Recall (verbal STM) involves immediate serial spoken recall of sequences of spoken digits. Backward Digit Recall (verbal WM) follows the same procedure except that participants attempt to recall the memory items in reverse sequence. The Dot Matrix test (visuo-spatial STM) involves serial recall of the locations of successive dots that appearing in an otherwise blank matrix. In Mr X (visuo-spatial WM), children make partial comparison judgment for a sequence of displays and then attempt serial recall of target spatial locations in the displays. *EFs.* The Switching and Planning subtests of the Delis-Kaplan Executive Function Scale (DKEFS, Delis, 2001) were administered. The planning subtest involved moving five disks of different sizes arranged on three pegs from a start position to an end state one disk at a time without placing any disk on a smaller disk. Total achievement scores were converted to scaled scores (M=10, SD=3). The switching subtest required children to connect letters and numbers in an increasing alternating sequence *(A-1-B-2*, etc.). Completion times were converted to scaled scores (M=10, SD=3). The Matrices subtest of nonverbal reasoning from the Wechsler Intelligence Scales for Children (Wechsler, 2014) was also administered. A pattern was presented with a missing piece, and children were required to select from a choice four below which piece completed the matrix. T-scores (M=50, SD=10) were calculated.

#### Speed

The Visual Scanning and Motor Speed subtests of the DKEFS were administered. Motor speed involved tracing a dotted line to connect circles as quickly as possible. The visual scanning test required children to cross out all the number threes on a response page. Completion times were converted to scaled scores as above.

### Learning measures

Standard scores were obtained for each of the following measures.

#### Vocabulary

The Peabody Picture Vocabulary Test-IV (Dunn & Dunn, 2007) assesses receptive vocabulary knowledge. Children are required to select one of four pictures showing the meaning of a spoken word.

#### Reading

The Single-Word Reading subtest of the Wechsler Individual Achievement Test II (WIAT II, Wechsler, 2005) assessed children’ s reading abilities. Children read a list of words aloud that were scored by the examiner. Responses were coded as correct if they were pronounced correctly and fluently.

#### Spelling

The Spelling subtest of the WIAT II n the provided a measure of spelling attainment. Children were asked to spell words spoken one at a time by the examiner.

#### Mathematics

The Number Operations subtest of the WIAT II was administered. This is an untimed test involving written mathematical problems that increase in difficulty.

### Behaviour measures

The Conners 3 Parent Rating Scale (Conners, 2008) assesses the presence of behavioural and cognitive problems related to ADHD. Parents or caregivers rate the frequency of 45 descriptions of problem behaviours. Scores on these items formed six subscales consisting of Inattention, Hyperactivity/ Impulsivity, Learning Problems, Executive Function, Aggression, and Peer Relations. The sum of raw scores on each subscale was converted to a T-score (M=50, SD=10). Data were incomplete for three children.

The Behavior Rating Inventory of Executive Function (BRIEF, Gioia, Isquith, Guy, & Kenworthy, 2000) questionnaire was completed by the parent of carer accompanying the child. The test contains 80 descriptions of problem behaviours relating to a wide range of executive functions, for which the adult rates the frequency. T-scores are derived for eight subscales: Inhibit, Shift, Emotional control, Initiate, Working memory, Planning, Organisation and Monitor. For one child, data were incomplete.

### Analysis

Following data checking and cleaning, the data were analysed using SPSSv22. Univariate outliers were identified for the cognitive and learning variables. Scores deviating from the sample mean by > 3.5 SDs were replaced with the highest / lowest allowable score (sample mean + / - 3.5 SDs). One multivariate outlier was detected using Mahalonbis *D*^*2*^ and was omitted from analysis.

The sample was divided into two sets of three subgroups based on i) learning profiles and ii) ADHD status. For each set of groups, group differences were investigated in separate MANOVAs on the learning, cognitive, and behaviour measures. Univariate F-tests were performed on significant group terms. Post hoc pairwise group comparisons were Bonferroni corrected for multiple comparisons on a family-wise basis (all dependent variables included in the analysis). Where the correction was applied, the reported *p* values are the original value multiplied by the number of post hoc comparisons within the family. Alpha was set at .05 for all analyses.

Exploratory factor analyses were performed on the cognitive and behaviour scores of this sample. Factors with eigenvalues > 1 are reported, with Varimax rotation to maximise differentiation. Separate MANOVAs were performed on the resulting factor scores with the sample grouped by either learning profile (low literacy, low maths, and low literacy and maths) and ADHD status (ADHD, ADHD under investigation, and no ADHD). Regression models of factor scores were conducted in order to test statistically possible interactions between groups. Finally, cognitive and behaviour factor scores were correlated with learning measures across the sample as a whole.

## Results

Mean reading, spelling and maths standard scores for the sample ranged between 80 and 85 (Table 1). Mean levels of performance close to age-appropriate levels were found for measures of speed of processing, matrix reasoning and planning. The measure showing the largest deficit in the sample was switching. All other scores fell in the low average range. Elevated levels of problem behaviours were observed across all behaviour subscales. Mean T-scores exceeded 60 (more than 1 *SD* from the population mean) on all measures, with scores indicating problems of clinical significance (70+) in inattention, hyperactivity/ impulsivity, learning problems, EFs, working memory and planning.

**Table 1.**
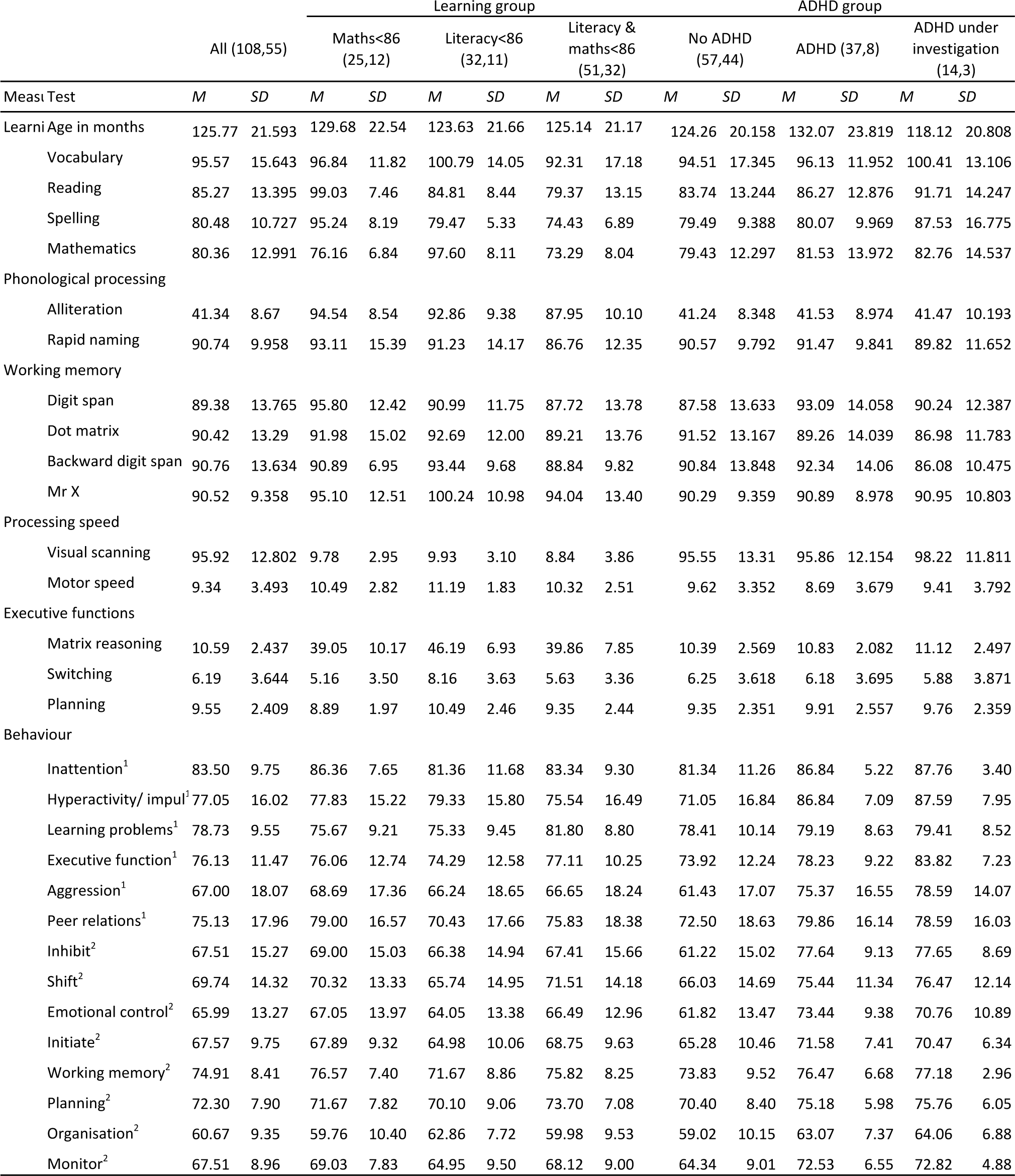
Descriptive statistics for the sample and by group (boys, girls)

### Learning and ADHD groups

Each child was grouped separately according to their learning profile: low literacy only (n=43, scores at or below 85 for either reading or spelling measures, or both), low maths only (n=37), and low literacy and maths (n=83). Descriptive statistics for the three learning subgroups as well as the same children grouped by ADHD status are shown in Table 2. The number of children in each combination of learning and ADHD groups is shown in S2. The association between the ADHD and learning subgroups was non-significant, χ^2^(4)=5.327, *p*>.05.

**Table 2.** Rotated component matrices for cognition and behavior

| Factor descriptors: | Cognition |  |  |
| --- | --- | --- | --- |
|  | F1<br>Executive/ | F2<br>Speed | F3<br>Phonological |
| Matrix | 0.684 | 0.105 | 0.203 |
| Alliteration | 0.175 | 0.040 | 0.668 |
| Rapid Naming | 0.026 | 0.690 | -0.068 |
| Digit Recall | -0.054 | -0.016 | 0.834 |
| Dot Matrix | 0.473 | 0.193 | 0.302 |
| Back Digit | 0.327 | 0.133 | 0.543 |
| Mr X | 0.637 | -0.098 | 0.286 |
| Visual Scan | 0.154 | 0.708 | 0.024 |
| Motor Speed | -0.083 | 0.712 | 0.120 |
| Switching | 0.378 | 0.526 | 0.147 |
| Planning | 0.727 | 0.086 | -0.200 |

| Factor descriptors: | Behaviour |  |  |
| --- | --- | --- | --- |
|  | F1<br>Hot EF | F2<br>ADHD | F3<br>Cool EF |
| Inattention | 0.180 | 0.809 | 0.030 |
| Hyper/ impuls | 0.599 | 0.557 | -0.173 |
| Learning problems | 0.069 | 0.038 | 0.653 |
| Executive functions | 0.095 | 0.787 | 0.289 |
| Aggression | 0.756 | 0.232 | 0.044 |
| Peer relations | 0.663 | -0.112 | 0.327 |
| Inhibit | 0.782 | 0.426 | 0.059 |
| Shift | 0.748 | 0.124 | 0.297 |
| Emotional control | 0.854 | 0.231 | 0.065 |
| Initiate | 0.480 | 0.313 | 0.562 |
| Working memory | 0.211 | 0.631 | 0.536 |
| Planning | 0.125 | 0.592 | 0.602 |
| Organisation | 0.228 | 0.664 | 0.073 |
| Monitor | 0.560 | 0.517 | 0.200 |

Correlations between measures of learning, cognition and behaviour are provided in S4. Separate exploratory factor analyses were performed on the cognitive and behaviour measures. The factor solutions are shown in Table 2.

#### Cognitive factors

Three cognitive factors accounted for 50.2% of the variance in scores. Factor 1 had highest loadings from measures associated with high executive loads and visuo-spatial aspects of cognition: matrix reasoning, measures of visuo-spatial STM and WM and of verbal WM, set switching and planning. The measures associated most strongly with the second factor either measured processing speed or were completed under time constraints: rapid naming, visual scanning, motor speed and switching. The three measures involving the manipulation or storage of phonological material (alliteration, forward and backward digit span) loaded strongly on the final factor. The factor descriptors are visuo-spatial (VS)/ executive, processing speed and phonological factors, respectively.

#### Behaviour factors

Corresponding analysis of the behaviour measures identified three factors that accounted for 65.6% of the variance. The first factor was linked with cool EFs (shifting, initiation of behaviour, and monitoring) as well as hot EFs (hyperactivity/ impulsivity, aggression, peer relations, inhibition, and emotional control). The second factor was most strongly associated with the symptoms of inattention and hyperactivity/ impulsivity that cooccur in ADHD, in addition to cool EFs (WM, planning, organisation, and monitoring). The third factor was associated exclusively with cool aspects of cognition and executive control: learning problems, initiation, WM and organisation. These factors are labelled hot EFs, ADHD symptoms, and cool EF, respectively.

#### Group analyses of factor scores

The top panel of Figure 1 shows the mean cognitive factor scores for the learning ability and ADHD groups. A MANOVA performed on these scores established a highly significant effect of learning group, *T* (6,314)=7.312, p<.001. The group term was significant on all three univariate F-tests: executive factor, p<.001; speed factor, p=.046, and phonological factor, p=.001. Pairwise post hoc comparisons showed significantly lower executive scores for the group with low maths and literacy than the children with low literacy alone, on the speed factor for the group with low literacy and maths than the low literacy group, and on the phonological factor for the group with low literacy and maths only than the group with low literacy alone. There was no significant effect of ADHD group on the cognitive factor scores, *T* (6, 314<1).

**Figure 1.**
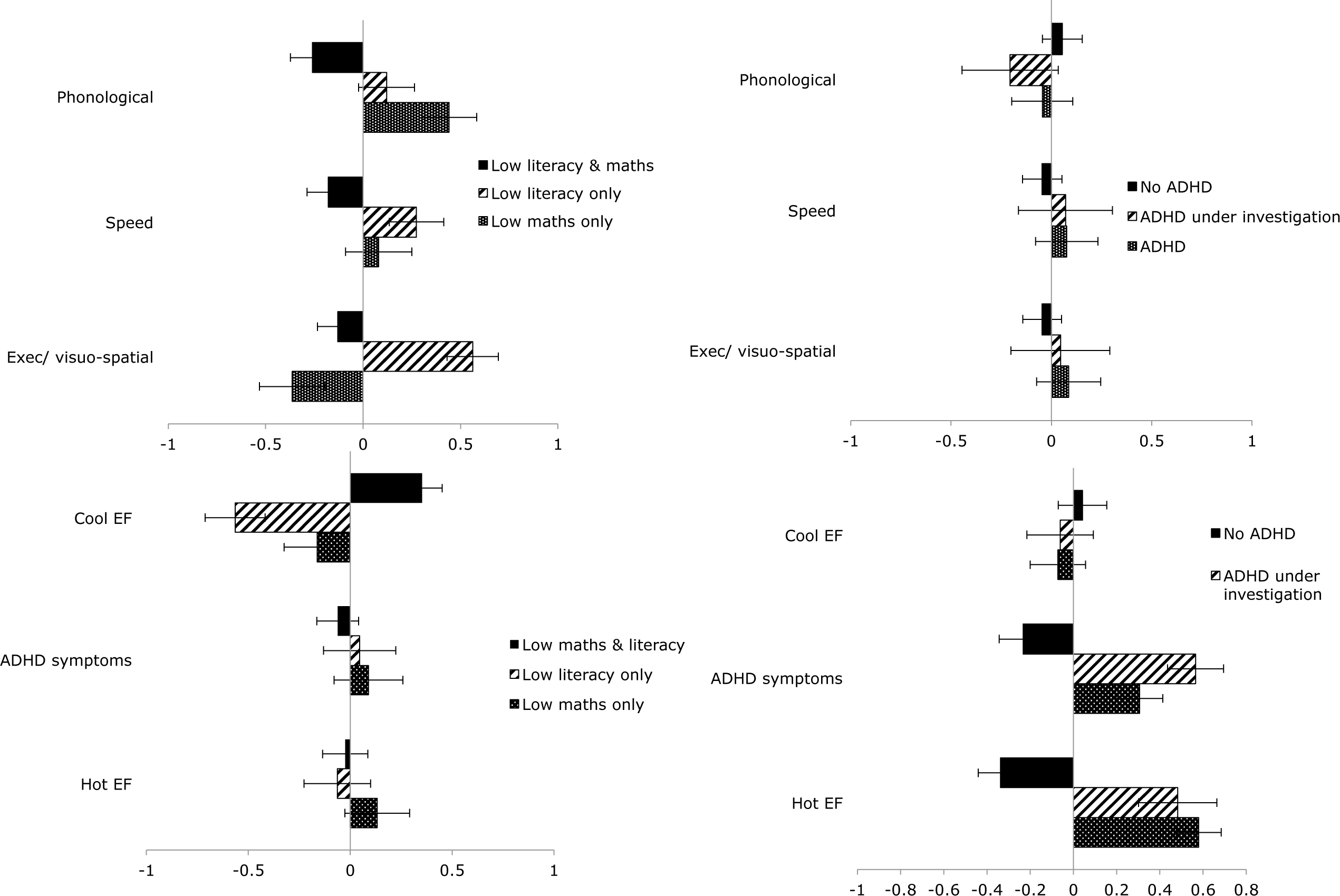
Top: mean cognitive factor scores (SEs) as a function of learning group (left) and ADHD group (right). Bottom: mean behaviour factor scores (Ses) as a function of learning group (left) and ADHD group (right).

The bottom panel of Figure 1 shows the behaviour factor scores derived as a function of learning ability and ADHD subgroups. A significant effect of learning ability group was found, *T* (6,304)=4.88, p<.001. By univariate F-tests, significant group differences were restricted to the cool EF factor, p<.001. Post hoc tests established the group low in both literacy and maths had significantly lower scores than those with either low literacy or low maths alone. In the corresponding analysis of behaviour factor scores by ADHD group, *T* (6,304)=9.987, p<.001, univariate group differences were significant only for the hot EF and ADHD symptom factors (p<.01). In both cases, post hoc comparisons established that this was due to higher factor scores in the group without ADHD than the ADHD or possible ADHD group (p<.01 in each case).

#### Regression analyses

The independence of ADHD status from learning subgroup was tested directly by comparing pairs of regression models performed on each of the six factor scores. The first model included three binary grouping factors: literacy (low, typical), maths (low, typical), and ADHD status (no ADHD, ADHD or probable ADHD). The second model added interaction terms for ADHD*literacy group and ADHD*maths group as a second block of independent variables. Model fit was then compared. If the relationships between cognition, behaviour and learning are independent of ADHD status as the previous analyses suggest, group interactions will not account for additional variance for any of the dimensions. The outcomes of the analyses are shown in S5.

For the executive and speed factors, the only significant predictor was mathematics group (p<.001 and *=.020*, respectively). Phonological scores were significantly predicted by both literacy group (p<.001) and less strongly by maths group (p=.022). Both the hot EF and ADHD symptoms factors were significantly associated with the ADHD group (p<.001 in both cases) but not with learning group. Cool EF scores were predicted both by literacy group (p=.007) and maths group (p<.001). Most importantly for the present purposes, the model fit did not significantly increase when the group interaction terms were added for any of the six factors. Changes in *R*^*2*^ were <.01 in each case.

### Whole-sample analysis

The classification of children into subgroups based on the conjunction of their maths and literacy scores does not align fully with a dimensional approach to learning abilities, as it treats subgroups as distinct categories. We therefore tested associations between learning and the cognitive and behaviour dimensions were tested across the entire sample. Correlations between each learning score (reading, spelling, and maths) and the six cognitive and behaviour factors are shown in Table 3. Reading and spelling abilities were significantly associated only with phonological skills (.379 and .279, respectively). Maths abilities were most strongly correlated with visuo-spatial executive skills (.413) but also shared links with both processing speed (.219) and phonological skills (.263). The correlation between reading and phonological skills remained significant when maths scores were partialled out (r=.375), as did the correlations between maths and each of the three cognitive dimensions (r=.409, .202 and .254 for the visuo-spatial/ executive, speed and phonological factors, respectively). The common links with phonological processing skills therefore could not be explained simply in terms of co-occurring deficits in both reading and maths. No significant links were found between learning abilities and either hot EF or ADHD symptoms of inattentive and hyperactivity/ impulsivity. The behavioural dimension of cool EF was weakly but significantly correlated with reading and spelling scores (r< .2 in both cases), and more strongly with maths scores (r=.318).

**Table 3.** Correlations between learning scores, cognitive and behaviour factors

| Measure | Learning |  |  |  | Cognition |  |  |
| --- | --- | --- | --- | --- | --- | --- | --- |
|  | Vocabulary | Reading | Spelling | Maths | Executive/<br>visuo-spatial | Speed | Phonological |
| Reading | .353** |  |  |  |  |  |  |
| Spelling | 0.154 | .735** |  |  |  |  |  |
| Maths | .266** | 0.111 | 0.102 |  |  |  |  |
| VS/ executive | .290** | 0.058 | -0.038 | .413** |  |  |  |
| Speed | 0.041 | 0.136 | 0.096 | .219** | 0 |  |  |
| Phonological | .418** | .379** | .297** | .263** | 0 | 0 |  |
| Hot behaviour | 0.006 | 0.133 | 0.078 | -0.02 | -0.093 | 0.115 | -0.003 |
| ADHD symptoms | 0.144 | 0.1 | 0.100 | 0.003 | 0.050 | -0.035 | -0.059 |
| Cool EF | -0.134 | -0.159* | -.174* | -.318** | -0.118 | -0.147 | -.205** |

## Discussion

This study examined the relationships between cognition, behaviour, learning and ADHD in children with problems in one or more areas of learning in a highly heterogeneous sample referred via health and education services. The primary aim was to discover whether the academic learning difficulties are associated with the same cognitive dimensions in children with and without ADHD. The simple answer is that they do. The cognitive skills of children who were struggling to learn with and without ADHD were indistinguishable. The same links between learning and separate cognitive dimensions were found for children diagnosed with ADHD, for those being investigated for possible ADHD, and for those without ADHD.

Three dimensions of cognitive abilities differentiated this sample of children. The first relates to skills in the processing and storage of phonological information, extending across measures of phonological awareness and verbal aspects of both STM and WM. The second dimension involves skills in tasks requiring storage and manipulation of nonverbal representations, extending across visuo-spatial aspects STM and WM and nonverbal problem solving. The final dimension is processing speed.

These dimensions had highly specific links with literacy and maths abilities, and these links were closely aligned with those reported in previous research on both typical and atypical populations. Literacy difficulties were most strongly limited by poor phonological skills, in line with the phonological deficit hypothesis (Bishop & Snowling, 2004; Melby-Lervag et al., 2012). Maths abilities, on the other hand, were associated most highly with the storage and high-level control of mental representations of nonverbal information such as patterns and locations in space. This fits well with evidence of selective links between mathematical abilities and visuo-spatial STM and WM as well as with other nonverbal and verbal executive (Moll, Göbel, Gooch, Landerl, & Snowling, 2016; Szucs, Devine, et al., 2013). Unique links were also found between maths abilities and phonological skills. This finding has been less commonly reported in previous studies, with low phonological abilities accompanying mathematical learning difficulties typically only when children also have reading problems (Moll et al., 2016; Szucs, Nobes, Devine, Gabriel, & Gebuis, 2013). Maths scores were linked with processing speed, but weakly.

The primary learning-related cognitive dimensions were therefore distinguished by informational domain: phonological and visuo-spatial. The links between these dimensions and learning also ran along domain-specific lines: phonological skills were most closely related to the acquisition of literacy, and the visuo-spatial executive skills to mathematical abilities. These cognitive dimensions cut across standard neuropsychological constructs of phonological awareness, STM, WM, and executive functions that are often employed as explanatory concepts in specific learning difficulties. Deficits in phonological processing and verbal aspects of STM co-exist in populations with reading and language impairments, suggesting a common deficit in handling phonology, the representational domain of language. This ability to represent verbal material in a phonological form appears to limit performance on the verbal measures in the CALM test battery. Of course, this does not mean that the cognitive composition of the tasks is identical. Other cognitive skills must also be required and these will be the source of task-specific variance likely to go undetected in dimension reduction methods.

The second domain-specific dimension tapped skills in storing and manipulating nonverbal material. Close links between visuo-spatial storage and more complex WM tasks have been frequently reported and interpreted as reflecting the relatively heavy burden on cognitive control imposed by forming and maintaining these representations (Alloway, Gathercole, & Pickering, 2006; Kane et al., 2004). This domain-specific cognitive control capacity appears to impose a significant constraint on mathematical learning in particular in the present sample.

In contrast, learning difficulties were unrelated to ADHD symptoms of inattention combined with hyperactive and impulsive behaviour, and to hot EFs involving emotional control. Both are common characteristics of children with ADHD (Sobanski et al., 2010). Across the sample as a whole, severity of learning difficulties was linked to problem behaviours such as failing to maintain attention across the course of an activity and having trouble remembering things, getting started on new activities and poor time management. These relate to cool EFs.

In summary, these findings favour a model in which learning difficulties originate in deficits in basic cognitive dimensions that include cool EFs, but ADHD arises from impairments in hot EFs. ADHD and learning difficulties appears to be consequence of coimpairments in two separate functional systems (Castellanos et al., 2005; Sonuga-Barke, 2002; Thorell, 2007; Zelazo & Müller, 2002). With comorbid disorders, the cognitive and behaviour symptoms summate but do not interact.

## Conclusion

This study successfully applies a dimensional approach to children with one or both of the two most common neurodevelopmental disorders - learning difficulties and ADHD. On the surface, children with the two kinds of disorder have little in common. Those with learning difficulties alone are far less hyperactive, impulsive and emotionally labile and experience fewer social problems than individuals with ADHD. Despite these differences, we have shown that many children with ADHD have the same deficits in basic phonological and visuo-spatial executive skills as children with learning problems alone. Moreover, they appear to give rise to the same learning problems. These findings have implications for both research and practice. First, the cognitive and behavioural characteristics of the mixed sample of children with learning difficulties either with ADHD or in isolation are more readily explained in terms of independent impairments in learning-critical cognitive dimensions and of the control of emotional and impulsive behaviour than of disorder-specific categorical diagnoses.Second, the dimensional profiles form the natural basis for the choice of the kinds of support and interventions necessary to meet the often complex needs of the individual child.

## Key points

- In a large sample of struggling learner, reading and maths achievements were related to three cognitive dimensions: phonological skills, speed of processing, and visuo-spatial/ executive skills.
- The links between cognition and learning were equivalent for children with and without ADHD
- Children with ADHD were distinguished only by their hyperactive and impulsive behaviours and other aspects of cognitive control related to affective value.
- The characteristics of children with learning difficulties either with ADHD or in isolation can be explained in terms of independent impairments in dimensions of cognition and behaviour.
- Dimensional profiles can form the natural basis for the choice of the kinds of support and interventions necessary to meet the needs of the individual child.

## Acknowledgments

This study was supported by the Medical Research Council and the University of Cambridge. Data collection was supported by research staff and postgraduate students at the MRC Cognition and Brain Sciences Unit. The authors wish to thank the many professionals working in children’ s services in the East of England for their support, and to the children and their families for giving up their time to visit the clinic.

---

**S1.**
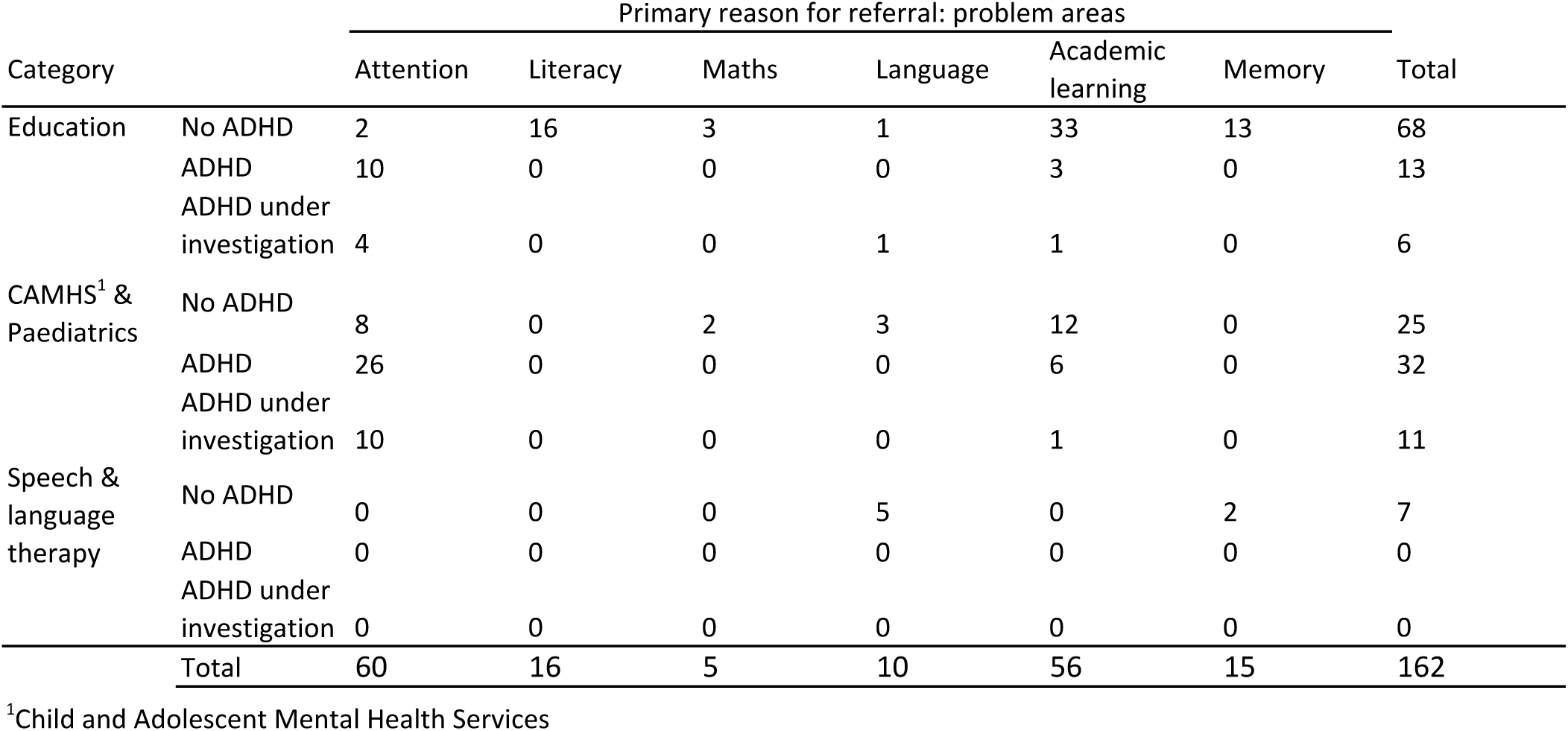
Number of children by referral route, ADHD status and primary reason for referral

**S2:**
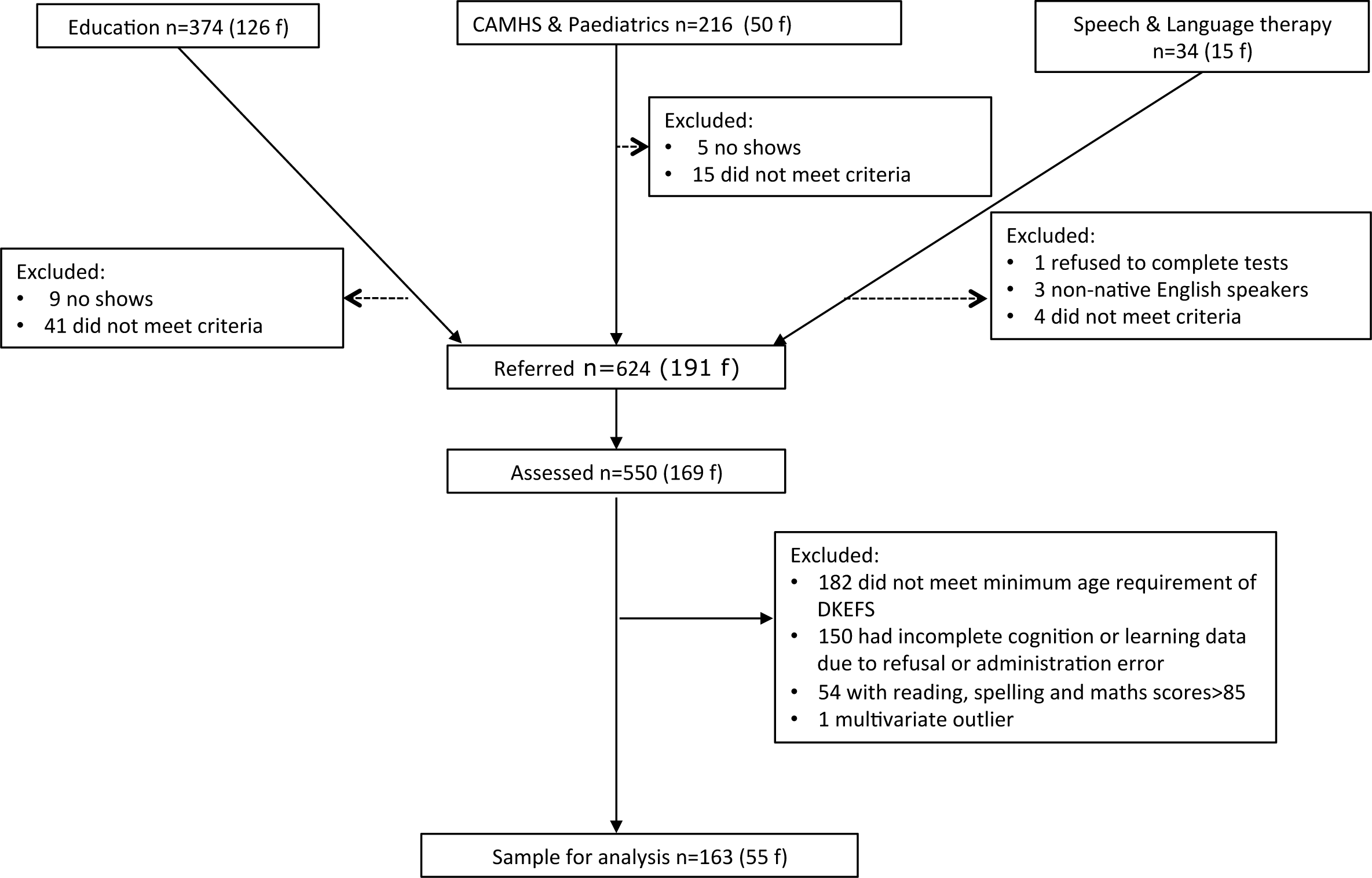
Recruitment flow chart showing recruitment routes and sources of exclusion to the data analysed

**S3.**
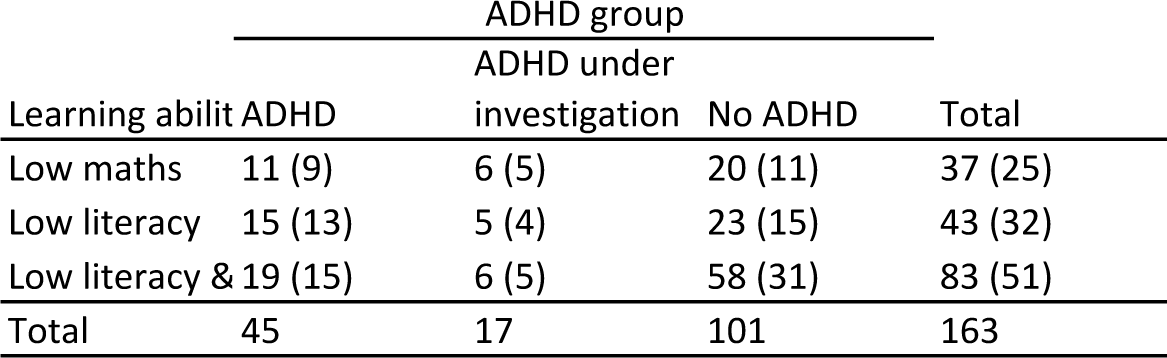
Numbers of children (m) as a function of learning ability and ADHD groups

**S4.**
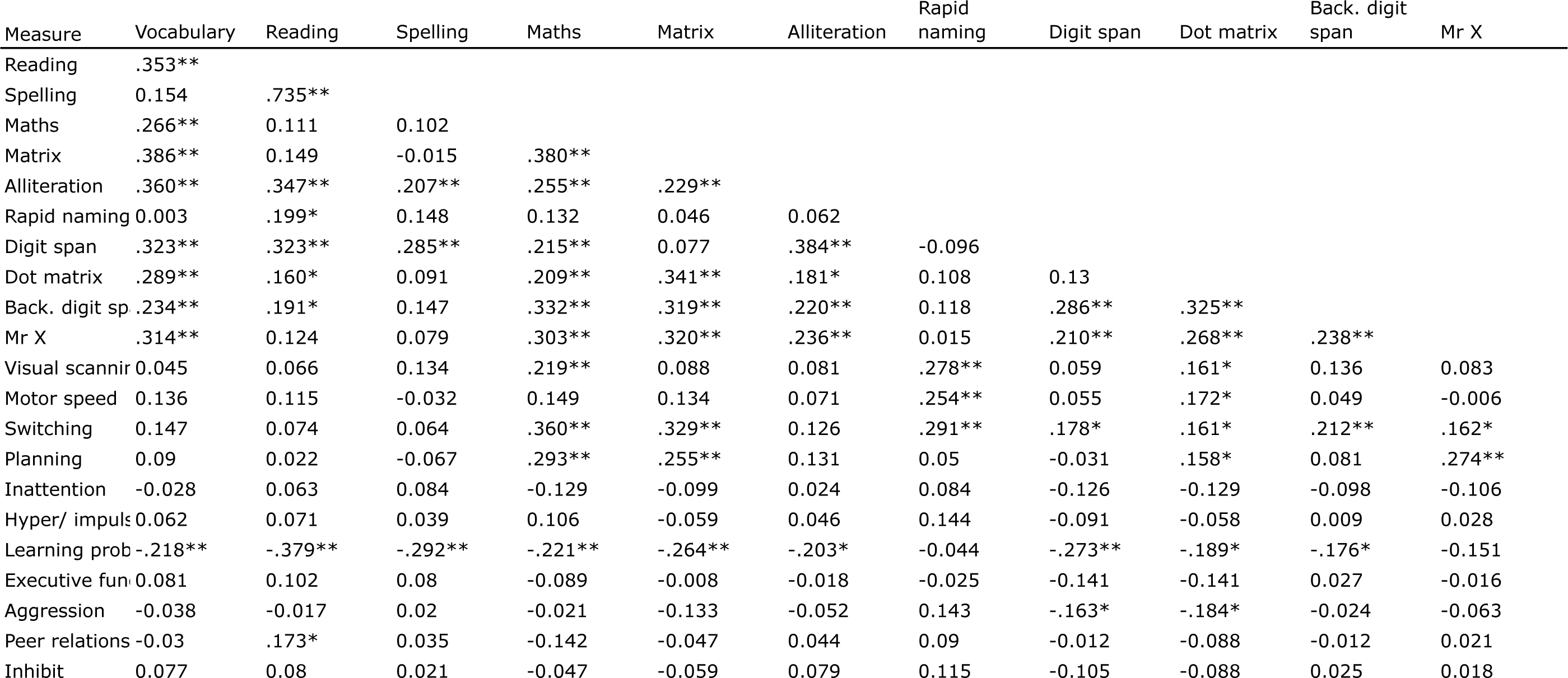
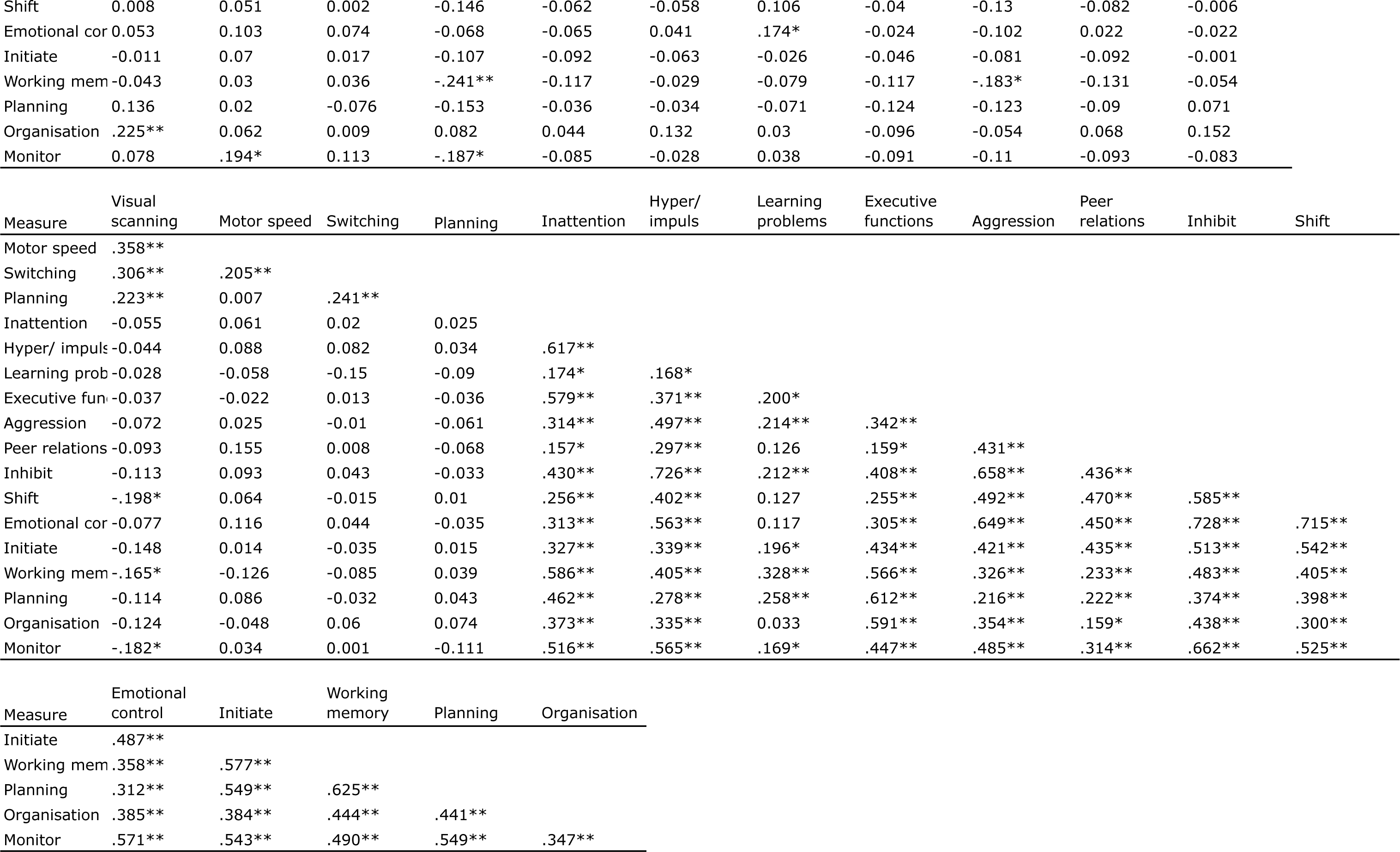
Correlations between learning, cognition and behaviour

**S5.**
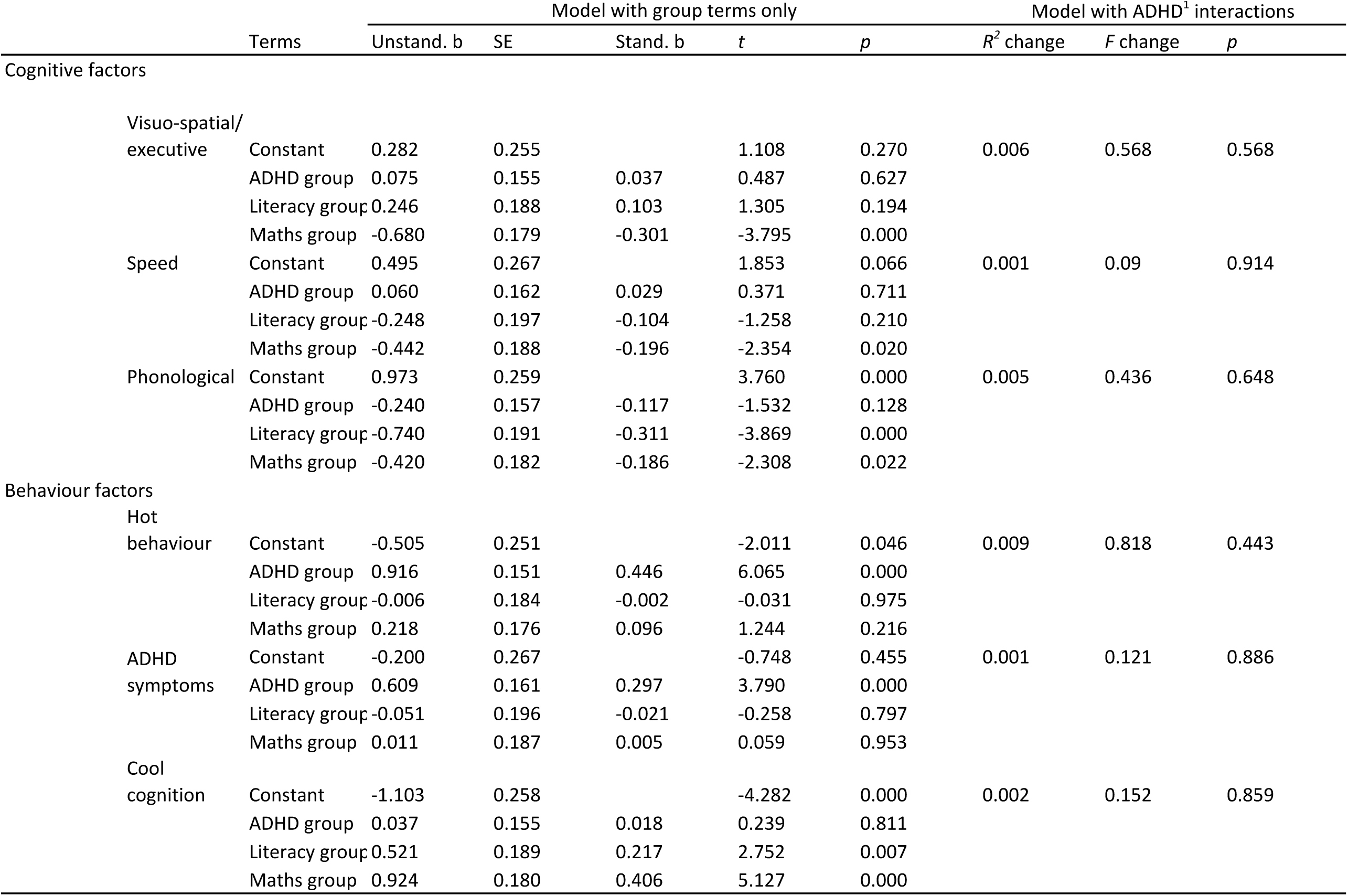
Regression models for cognitive and behaviour factors with and without ADHD by learning group interactions

## References

Alloway, T. P. (2007). Automated Working: Memory Assessment: Manual: Pearson.

Alloway, T. P., Gathercole, S. E., & Pickering, S. J. (2006). Verbal and visuospatial short-term and working memory in children: are they separable? Child Development, 77(6), 1698–1716. doi:10.1111/j.1467-8624.2006.00968.x

APA. (2013). Diagnostic and statistical manual of mental disorders (DSM-5®): American Psychiatric Pub.

Bishop, D. V., & Snowling, M. J. (2004). Developmental dyslexia and specific language impairment: same or different? Psychological bulletin, 130(6), 858–886. doi:10.1037/0033-2909.130.6.858

Bull, R., & Scerif, G. (2001). Executive functioning as a predictor of children’ s mathematics ability: inhibition, switching, and working memory. Developmental neuropsychology, 19(3), 273–293. doi:10.1207/S15326942DN1903_3

Casey, B., Oliveri, M. E., & Insel, T. (2014). A neurodevelopmental perspective on the research domain criteria (RDoC) framework. Biological Psychiatry, 76(5), 350–353.

Castellanos, F. X., Sonuga-Barke, E. J., Scheres, A., Di Martino, A., Hyde, C., & Walters, J. R. (2005). Varieties of attention-deficit/hyperactivity disorder-related intra-individual variability. Biological Psychiatry, 57(11), 1416–1423.

Cheung, C. H. M., Wood, A. C., Paloyelis, Y., Arias-Vasquez, A., Buitelaar, J. K., Franke, B., … Kuntsi, J. (2012). Aetiology for the covariation between combined type ADHD and reading difficulties in a family study: the role of IQ. Journal of Child Psychology and Psychiatry, 53(8), 864–873. doi:10.1111/j.1469-7610.2012.02527.

x Conners, C. K. (2008). Conners Parent Rating Scale 3rd edition: Pearson Publishing.

Cuthbert, B. N., & Insel, T. R. (2013). Toward the future of psychiatric diagnosis: the seven pillars of RDoC. BMC medicine, 11(1), 126.

Delis, D. (2001). Delis-Kaplan executive function scale (D-KEFS). San Antonio: The Psychological Corporation.

Dunn, L. M., & Dunn, D. M. (2007). PPVT-4: Peabody picture vocabulary test: Pearson Assessments.

Gathercole, S. E. (2006). Nonword repetition and word learning: The nature of the relationship. Applied Psycholinguistics, 27(04), 513–543.

Gioia, G. A., Isquith, P. K., Guy, S. C., & Kenworthy, L. (2000). Behavior rating inventory of executive function: BRIEF: Psychological Assessment Resources Odessa, FL.

Gremillion, M. L., & Martel, M. M. (2012). Semantic language as a mechanism explaining the association between ADHD symptoms and reading and mathematics underachievement. Journal of abnormal child psychology, 40(8), 1339–1349.

Holmes, J., Hilton, K. A., Place, M., Alloway, T. P., Elliott, J. G., & Gathercole, S. E. (2014). Children with low working memory and children with ADHD: same or different? Frontiers in human neuroscience, 8, 976. doi:10.3389/fnhum.2014.00976

Kane, M. J., Hambrick, D. Z., Tuholski, S. W., Wilhelm, O., Payne, T. W., & Engle, R. W. (2004). The generality of working memory capacity: a latent-variable approach to verbal and visuospatial memory span and reasoning. Journal of experimental psychology. General, 133(2), 189–217. doi:10.1037/0096-3445.133.2.189

McGrath, L. M., Pennington, B. F., Shanahan, M. A., Santerre-Lemmon, L. E., Barnard, H. D., Willcutt, E. G., … Olson, R. K. (2011). A multiple deficit model of reading disability and attention-deficit/hyperactivity disorder: Searching for shared cognitive deficits. Journal of Child Psychology and Psychiatry, 52(5), 547–557.

Melby-Lervag, M., Lyster, S.-A. H., & Hulme, C. (2012). Phonological skills and their role in learning to read: a meta-analytic review: American Psychological Association.

Moll, K., Göbel, S. M., Gooch, D., Landerl, K., & Snowling, M. J. (2016). Cognitive risk factors for specific learning disorder processing speed, temporal processing, and working memory. Journal of learning disabilities, 49(3), 272–281.

Muter, V., Hulme, C., & Snowling, M. J. (1997). The phonological abilities test: The Psychological Corporation.

Ramus, F., Marshall, C. R., Rosen, S., & van der Lely, H. K. (2013). Phonological deficits in specific language impairment and developmental dyslexia: towards a multidimensional model. Brain : a journal of neurology, 136(Pt 2), 630–645. doi:10.1093/brain/aws356

Sobanski, E., Banaschewski, T., Asherson, P., Buitelaar, J., Chen, W., Franke, B., … Faraone, S. V. (2010). Emotional lability in children and adolescents with attention deficit/hyperactivity disorder (ADHD): clinical correlates and familial prevalence. Journal of Child Psychology and Psychiatry, 51(8), 915–923. doi:10.1111/j.1469-7610.2010.02217.x

Solanto, M. V., Arnsten, A. F. T., & Castellanos, F. X. (2001). Stimulant drugs and ADHD: Basic and clinical neuroscience: Oxford University Press, USA.

Sonuga-Barke, E. J. S. (2002). Psychological heterogeneity in AD/HD—a dual pathway model of behaviour and cognition. Behavioural brain research, 130(1), 29–36.

Sonuga-Barke, E. J. S., Bitsakou, P., & Thompson, M. (2010). Beyond the dual pathway model: evidence for the dissociation of timing, inhibitory, and delay-related impairments in attention-deficit/hyperactivity disorder. Journal of the American Academy of Child & Adolescent Psychiatry, 49(4), 345–355.

Sonuga-Barke, E. J. S., & Coghill, D. (2014). Editorial Perspective: Laying the foundations for next generation models of ADHD neuropsychology. Journal of Child Psychology and Psychiatry, 55(11), 1215–1217. doi:10.1111/jcpp.12341

Swanson, H. L., & Sachse-Lee, C. (2001). A subgroup analysis of working memory in children with reading disabilities: domain-general or domain-specific deficiency? Journal of learning disabilities, 34(3), 249–263.

Szucs, D., Devine, A., Soltesz, F., Nobes, A., & Gabriel, F. (2013). Developmental dyscalculia is related to visuo-spatial memory and inhibition impairment. Cortex; a journal devoted to the study of the nervous system and behavior, 49(10), 2674–2688. doi:10.1016/j.cortex.2013.06.007

Szucs, D., Nobes, A., Devine, A., Gabriel, F. C., & Gebuis, T. (2013). Visual stimulus parameters seriously compromise the measurement of approximate number system acuity and comparative effects between adults and children. Frontiers in psychology, 4, 444. doi:10.3389/fpsyg.2013.00444

Thorell, L. B. (2007). Do delay aversion and executive function deficits make distinct contributions to the functional impact of ADHD symptoms? A study of early academic skill deficits. Journal of Child Psychology and Psychiatry, 48(11), 1061–1070.

Wechsler, D. (2005). Weschler Indivisual Achievement Test II: Pearson Publishing.

Wechsler, D. (2014). Wechsler Intelligence Scale for Children V: NCS Pearson, Incorporated.

Willcutt, E. G., Betjemann, R. S., McGrath, L. M., Chhabildas, N. A., Olson, R. K., DeFries, J. C., & Pennington, B. F. (2010). Etiology and neuropsychology of comorbidity between RD and ADHD: The case for multiple-deficit models. Cortex; a journal devoted to the study of the nervous system and behavior, 46(10), 1345–1361.

Zelazo, P. D., & Müller, U. (2002). Executive function in typical and atypical development.

Zhao, Y., & Castellanos, F. X. (2016). Annual Research Review: Discovery science strategies in studies of the pathophysiology of child and adolescent psychiatric disorders-promises and limitations. Journal of Child Psychology and Psychiatry, 57(3), 421–439.

